# Grain aphids (*Sitobion avenae*) with knockdown resistance (kdr) to insecticide exhibit fitness trade-offs, including increased vulnerability to the natural enemy *Aphidius ervi.*

**DOI:** 10.1101/2020.03.04.976332

**Authors:** Damon Little, Gaynor Malloch, Louise McNamara, Gail E. Jackson

## Abstract

The development of insecticide-resistance mechanisms in aphids has been associated with inhibitory, pleiotropic fitness costs. Such fitness costs have not yet been examined in the UK’s most damaging cereal aphid, *Sitobion avenae* (grain aphid) (Hemiptera: Aphididae). This study aimed to evaluate the fitness trade-offs of the insecticide-resistant *S. avenae* clone versus an insecticide-susceptible *S. avenae* clone. Additionally, the parasitoid, *Aphidius ervi* (Hymenoptera: Braconidae), was introduced to examine its potential as a biological control agent. This study found that insecticide-resistant clones had significantly lower population growth and individual relative growth rate. Furthermore, insecticide-resistant clones suffered from a significantly greater rate of parasitisation (mummification) compared to their insecticide-susceptible counterparts. The successfulness of the parasitoid as a biological control agent could prevent the spread of the insecticide-resistant genotype. However, for this to be possible, insecticide spraying regimes need to be moderated, and habitat modification and parasitoid manipulation must be considered.

## Introduction

The evolution of organisms occurs via genetic variation and selection imposed by many abiotic and biotic environmental factors. Each factor can exert an opposing selection pressure, resulting in the variation of optimal levels of defence or immunity depending on the environmental conditions. The establishment of trade-offs occur when opposing selection pressures cause the defence/immunity level to be lower than the maximum (Boivin *et al*., 2003). In areas where selection pressures vary over time, the balance between trade-offs can shift, which may lead to the optimal defence/immunity level changing. Environmental fluctuations can lead to organisms mutating to better suit their new environment; however, these mutations can be limited if they incur pleiotropic fitness costs which affect physiological or behavioural traits (Boivin *et al*., 2003).

In response to strong, unambiguous selection pressures caused by intense, widespread agricultural activity, some pests have developed adaptive traits including pesticide resistance. A mechanism known as ‘knockdown resistance’ (kdr) has allowed cross-resistance in pests to DDT and pyrethroids. This mechanism is characterised by a reduction in the sensitivity of the nervous system caused by a single amino acid substitution (L1014F) in the insect’s voltage gated sodium channel gene (Sawicki, 1985; Foster *et al*. 2014). Intuitively, kdr resistant individuals should experience fitness costs in areas where there is no insecticide pressure compared to susceptible ones. If this was not the case, the frequency of resistant alleles would be higher prior to exposure to pesticides (Crow, 1957). Therefore, the resistant genes are likely to have deleterious pleiotropic costs, which have constrained the adaptive trait (Roush and McKenzie, 1987). In support of this theory, there is growing evidence detailing the maladaptive side-effects of fitness changes on other seemingly unrelated traits (Foster et al. 2003; Foster et al. 2007).

During the late summer of 2011, growers in England began reporting that *Sitobion avenae* (grain aphid) (Hemiptera: Aphididae) were becoming less susceptible to pyrethroids sprayed on cereal crops. *S. avenae* is one of the most damaging cereal aphids in Western Europe, feeding on all cereals including barley, wheat and rice (Van Emden and Harrington, 2017). *S. avenae* show a strong preference for the ear of cereals, which generally stay physiologically active for longer than the leaf. This allows *S. avenae* to maintain itself for considerably longer than other aphid species (Dean and Luuring, 1970). Foster *et al*. (2014) identified that the kdr mechanism had resulted in clonal variation in the *S. avenae* sample with resistant clones exhibiting a 40-fold Resistance Factor. Currently, most studies investigating fitness trade-offs caused by insecticide-resistance have involved *Myzus persicae* (peach-potato aphid). There are no studies into the effect of the kdr mechanism on *S. avenae* and the maladaptive fitness traits that may be incurred. Existing literature suggests that the kdr-resistant *S. avenae* clone may have invested in the kdr mutation at the cost of pleiotropic performance traits. Malloch *et al*. (2016) show that the frequency of the kdr resistant *Sitobion avenae* clone in UK suction trap catches has stabilised at around 30%, which provides further evidence for the likelihood of some fitness costs associated with the kdr mutation. Currently the kdr mechanism is heterozygous (kdr-SR), but if homozygous resistance (kdr-RR) were to evolve, the levels of resistance would be expected to further increase (Foster *et al*. 2014).

With increased resistance to pesticides, it has become imperative to develop other pest management techniques, such as, exploiting and manipulating the natural enemies of pests to act as a biological control (Huffaker, 2012). The use of natural enemies to supress specific pest organisms has evolved into an important facet of integrated pest management (IPM) (e.g. Hunter, 1909, Room *et al*. 1981). The effectiveness of natural enemies as a biological control depends on several characteristics. These include high reproductive potential, a short development time in relation to prey and a high level of prey specificity (Debach and Rosen, 1991). Such characteristics are exemplified in the parasitoid Diptera and Hymenoptera. Adult females belonging to these orders are generally highly fecund, develop inside their prey making generation time similar to that of the host and only specialise in attacking a small number of prey species (Debach and Rosen, 1991). With over 400 species recorded (Starý *et al*. 1988), the use of aphid-specific parasitoids (Hymenoptera: Braconidae) in controlling aphid populations has been well documented in various cropping systems (Chambers *et al*. 1986; Starý *et al*. 1988). *Aphidius ervi*, used in this study is a solitary endophagous parasitoid, with an overall time from oviposition to wasp emergence of 14 ± 3 days (Thiboldeaux, 1986; Ives et al. 1999). To locate hosts, *A. ervi* use chemical cues such as aggregation and sex pheromones, and plant volatiles (Godfray, 1994). After locating the aphid, female *A. ervi* rapidly attempt to parasitise it by penetrating its exoskeleton with an ovipositor (Starý, 1988).

The present study was designed to determine if the kdr-resistant *S. avenae* clone has developed any maladaptive behavioural or physiological characteristics because of the kdr mechanism. The study compared and assessed differences in performance traits in the kdr-resistant and kdr-susceptible clones. It was hypothesised that the kdr-susceptible clones would have a significantly greater aphid population growth rate and individual relative growth rate than the kdr-resistant clones. Additionally, following the introduction of the parasitoid, *Aphidius ervi* (Hymenoptera: Braconidae), it was hypothesised that the kdr-susceptible clone would be able to deter the parasitic wasp more successfully than the kdr-resistant clone. Furthermore, it was hypothesised that in comparison with the kdr-susceptible clone, a greater proportion of kdr-resistant clone would be parasitised, and the parasitoid emergence rate would be greater in the kdr-resistant colonies.

## Materials and Methods

### Study Species

Barley (*Hordeum vulgare* cv. Sienna) was used as the host plant. Four barley seeds were planted in to each of 24 2 L pots containing Levington M3 High Nutrient Compost (Everris, Ipswich, UK). Plants were grown in a glasshouse at 21 ± 2 °C, under a 16:8 light:dark photoperiod and watered twice weekly throughout the study. After 21 days plants were thinned to leave one plant per pot, which were grown on for a further 40 days until they reached GS23 (AHDB Cereal growth stages).

Two clonal lines of the grain aphid, *Sitobion avenae*, were sourced from long-term colonies reared by the James Hutton Institute in Dundee: (1) homozygous fully insecticide susceptible of SA12A lineage (kdr-SS) and (2) kdr heterozygous insecticide resistant of SA3 lineage (kdr-SR). Aphid colonies were reared on three barley plants in separate mesh cages (50 cm by 50 cm by 50 cm) in an insectary (17 ± 3 °C; 65 ± 5 % RH; LD 16:8 h, 150 μmol m^-2^ s^-1^). Plants were replenished each week. Clonal integrity was verified at the beginning and end of the experiment through DNA genotyping. DNA was extracted from single adult *S. avenae* using a sodium hydroxide method described in Malloch et al. (2006). Five microsatellite loci were examined: Sm10, Sm12, Sm17, SaΣ4, and S16b using published primer pair sequences (Simon et al. 1999, Wilson et al. 2004). PCR was carried out in 8 μl volumes using Illustra™ Ready to Go PCR beads (GE Healthcare). When the bead is reconstituted the concentration of each dNTP is 200 μM in 10 mM Tris HCl, 50 mM KCl and 1.5 mM MgCl_2_. Each bead contains 2.5 units of Taq polymerase. PCR was carried out on a Techne 5 Prime /02 thermal cycler using the Touchdown programme described in Sloane et al. (2001). Genotyping was carried out on an ABI 3730 DNA analyser and the results interpreted using GeneMapper software (Applied Biosystems 2005). Genotypes were assigned using a reference data set for the SA12A and SA3 colonies held at the James Hutton Institute.

The aphid parasitoid *Aphidius ervi*, was acquired as mummies (Fargro Ltd., West Sussex) and used immediately upon receipt.

### Experimental Setup

The experiment was conducted in Scotland’s Rural College (SRUC) insectary (17 ± 3 °C; 65 ± 5 % RH; LD 16:8 h, 150 μmol m^-2^ s^-1^) at the King’s Buildings campus at the University of Edinburgh between the 11^th^ February 2019 and the 5^th^ April 2019.

61 days after planting (at GS23), 24 barley plants were randomly assigned to eight mesh chambers (50 cm by 50 cm by 50 cm), with three plants per chamber. The kdr-SS and kdr-SR clones were randomly allocated to each chamber so that there were four chambers of each genotype. Each plant was inoculated with six apterous adult aphids. They were distributed evenly between the first and second longest tiller of the plant (three aphids per tiller).

### Aphid Performance Traits

#### Tiller-level aphid abundance

Aphid counts were conducted twice a week for five weeks on the first and second longest tiller of each plant. All aphids from the base of the tiller to the ear were counted. This acted as a proxy of aphid population growth.

#### Aphid Relative Growth Rate (RGR)

RGR was calculated for one aphid per plant. A clip cage (Noble, 1958) was placed over a healthy apterous adult aphid. After 24 h the clip cage was removed and the adult aphid and all but one of the nymphs were removed. The clip cage was then replaced over the nymph, and after a further 48 hours, it was weighed using a Mettler Toledo XP6 Analytical Balance (Mettler Toledo Ltd., Leicester, UK). After weighing, it was transferred back to the barley leaf and covered by the clip cage again. Exactly 72 h later, the nymph was reweighed, and RGR calculated using van Emden and Bashford’s (1969) formula:

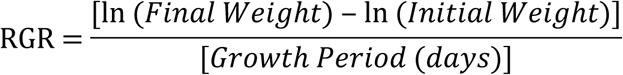

The RGR values were then averaged for each genotype to create a Mean Relative Growth Rate (MRGR). On the occasion that the aphid final weight was less than the initial weight, the data were discarded due to the assumption that the aphid had been damaged (Jackson, 1995).

#### Parasitoid-aphid interactions

Aphid behaviour responses initiated by parasitoid wasps were observed under a binocular microscope. A 5 cm length of barley leaf with one apterous adult aphid attached was placed inside a Perspex petri dish along with one female wasp. Following first physical contact between the aphid and wasp, aphid behavioural responses were recorded for one minute. First contact occurred when the parasitoid walked over the aphid, or touched it with its ovipositor or antennae (Foster *et al*. 2007). During this time, the ‘warding behaviour’ recorded as the number of kicks and drops were counted. A kick was defined as the aphid moving its body vigorously whilst kicking its hind legs in the direction of the wasp (Dixon, 1988). A drop was recorded when the aphid did a short jump away from the feeding site and the wasp. This normally resulted in the aphid detaching itself from the leaf (Villagra *et al*. 2002). A new wasp, aphid, barley leaf and Perspex petri dish was used for each observation to avoid pseudoreplication.

#### *Mummification of* Sitobion avenae *clones*

Thirty one days after aphids were placed on the experimental plants (92 days after planting), thirty-five *A. ervi* wasp mummies were placed into each of eight Perspex petri dishes, one of which was added to the centre of each chamber. The emergent wasps were left in the chambers for 21 days to parasitise the aphids. Each barley plant was then harvested along with its aphid population, and the number of new mummies per plant counted, removed and placed into sealed petri dishes. Each plant was then bagged and placed into a freezer (−20 °C) for three days. The number of aphids per plant was then counted and the proportion of mummified aphids to the total number of aphids calculated for each plant.

#### Aphidius ervi *emergence success*

The Petri dishes containing the collected mummies remained in the insectary for seven days to allow the parasitoids to emerge freely, in accordance with development times determined by Ives *et al*. (1999). The number of hatched mummies was then counted. Each mummy was examined for an emergence hole, and the proportion hatched to unhatched represented parasitoid emergence success.

### Data Analysis

R version 3.5.1 statistical software was used to conduct all statistical analyses (R Core Team, 2018). Since aphid count data was not normally distributed, the natural log was taken and used throughout. All data were checked for normal distribution using a Shapiro-Wilk test. After this assumption was met, a Bartlett test was carried out on categorical data to ensure there was equal variance across all samples (homoscedasticity). All the data also met this assumption.

A Mixed Effects Model was used to examine the relationship between days and number of aphids. Days were included as a random effect because they were not independent of one another, with days closer to each other being more likely to be interrelated than ones further apart. It was assumed that data collected from plants within a chamber may be correlated, therefore, plants were nested within chambers and included as an additional random effect. This allowed for an increased sample size and prevented the need to aggregate data through averaging.

The natural log of aphid counts were averaged for each chamber for each day measurements were taken. A single-factor analysis of variance (ANOVA) assessed whether there was a significant difference between the kdr-SS and kdr-SR aphid counts performed on each day. A single-factor ANOVA was also used to determine differences between genotypes for MRGR, proportion of mummified aphids and emergence success. The interaction data was normally distributed count data. Therefore, a generalised linear model with Poisson distribution was used to assess the effect of genotype.

Graphs were made using Microsoft Excel 2016 or SigmaPlot 13.0. When constructing Figure 1, raw data were used rather than logged values. This showed variation and error more clearly.

**Figure 1:**
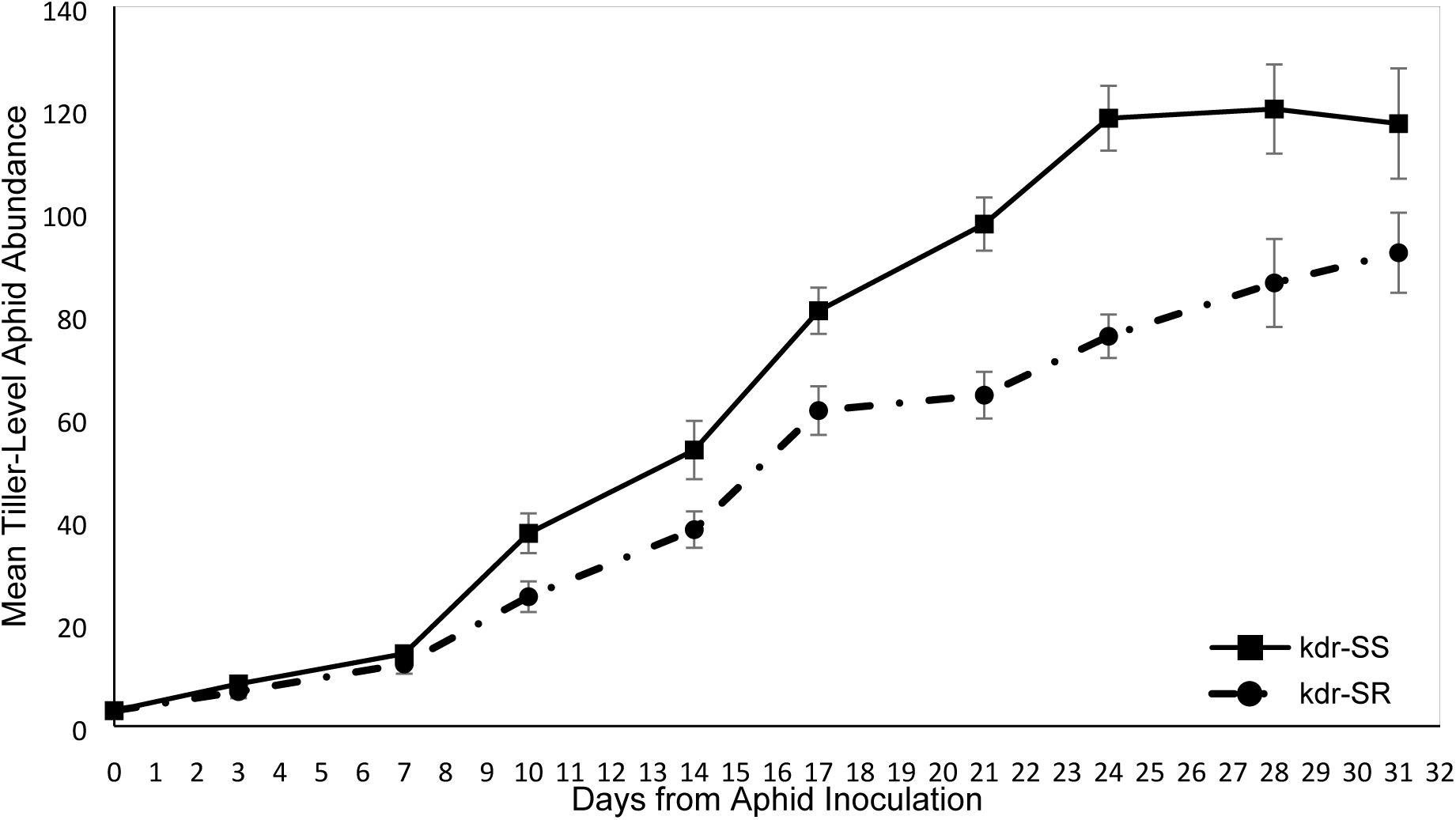
Mean tiller-level aphid abundance over time. Error bars represent ± SE for each category for each day of measurement. (n = 24)

### Results

#### Tiller-level aphid abundance

Figure 1 shows that the kdr-SS clone had greater tiller-level abundance throughout the experiment than the kdr-SR clone. From day 7 onwards this was statistically significantly different. The kdr-SR clone increased, on average, by 3.2 aphids per day, whereas the kdr-SS clone increased by an average of 4.4 aphids per day. The significant difference in abundance between clones continued until day 28, by which point the kdr-SS abundance had reached a plateau of approximately 119 ± 2 individuals. The kdr-SR population was still increasing at the end of the 31 day period.

#### Mean Relative Growth Rate

Figure 2 illustrates that the Mean Relative Growth Rate (MRGR) of individual kdr-SR aphids was significantly lower than that of individual kdr-SS clone (F_1,14_ = 4.8, p < 0.0466). Sample size varied between genotype due to aphid damage or mortality.

**Figure 2.**
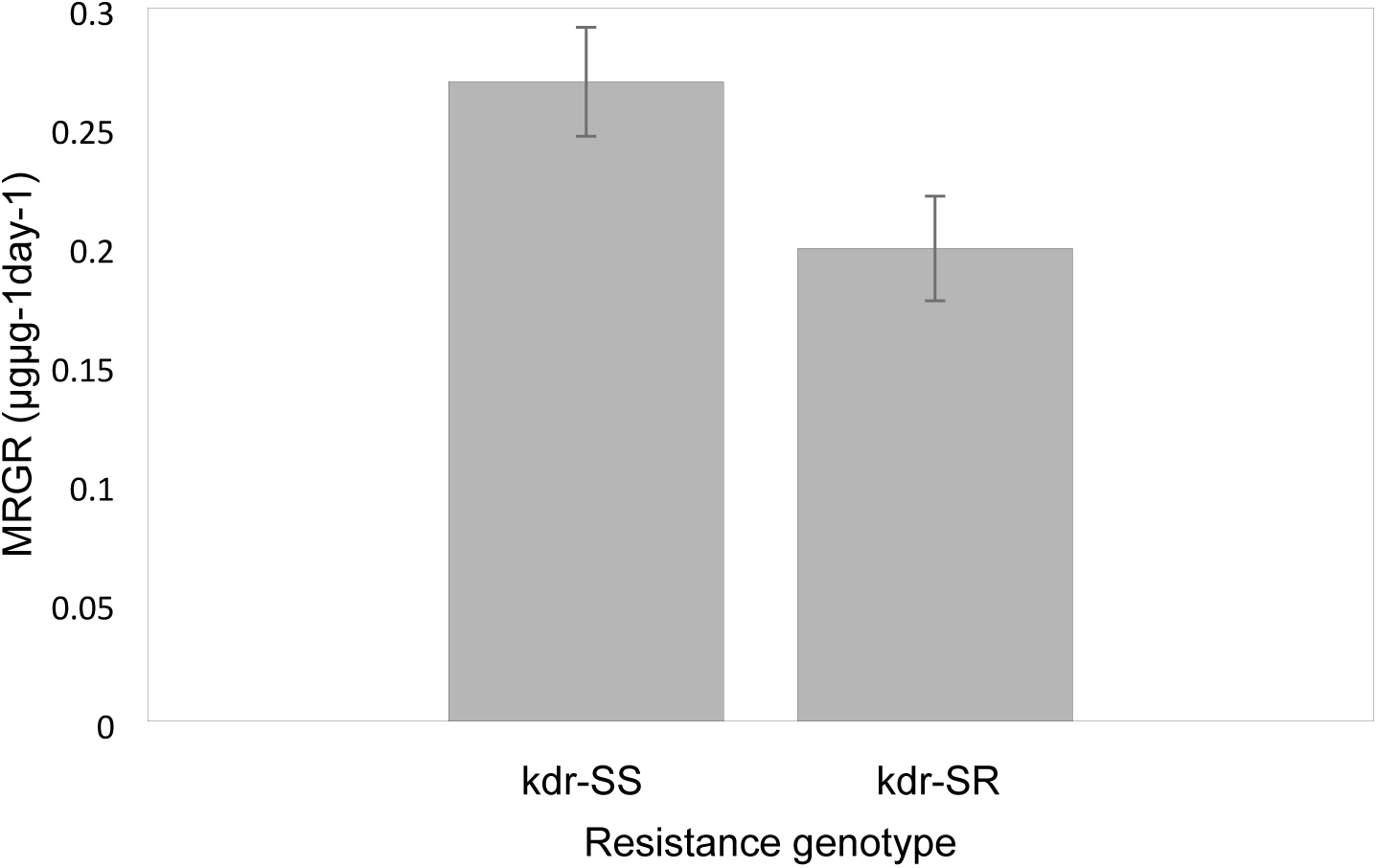
Mean Relative Growth Rate (MRGR) of the kdr-SS and kdr-SR clones. Error bars represent ± individual SE for each category. Aphids that were damaged were not included in the analysis. (kdr-SS *n* = 7, kdr-SR *n* = 9).

#### Parasitoid-aphid interaction

Figure 3a illustrates that there was no difference in the mean number of ‘warding behaviours’ between the kdr-SR and kdr-SS clones when attacked by a wasp. (GLM - *z*-value = 0.29, *df* = 39, *p* < 0.77). The mean number of kicks per minute was 10.5 ± 0.5 for both genotypes.

**Figure 3.**
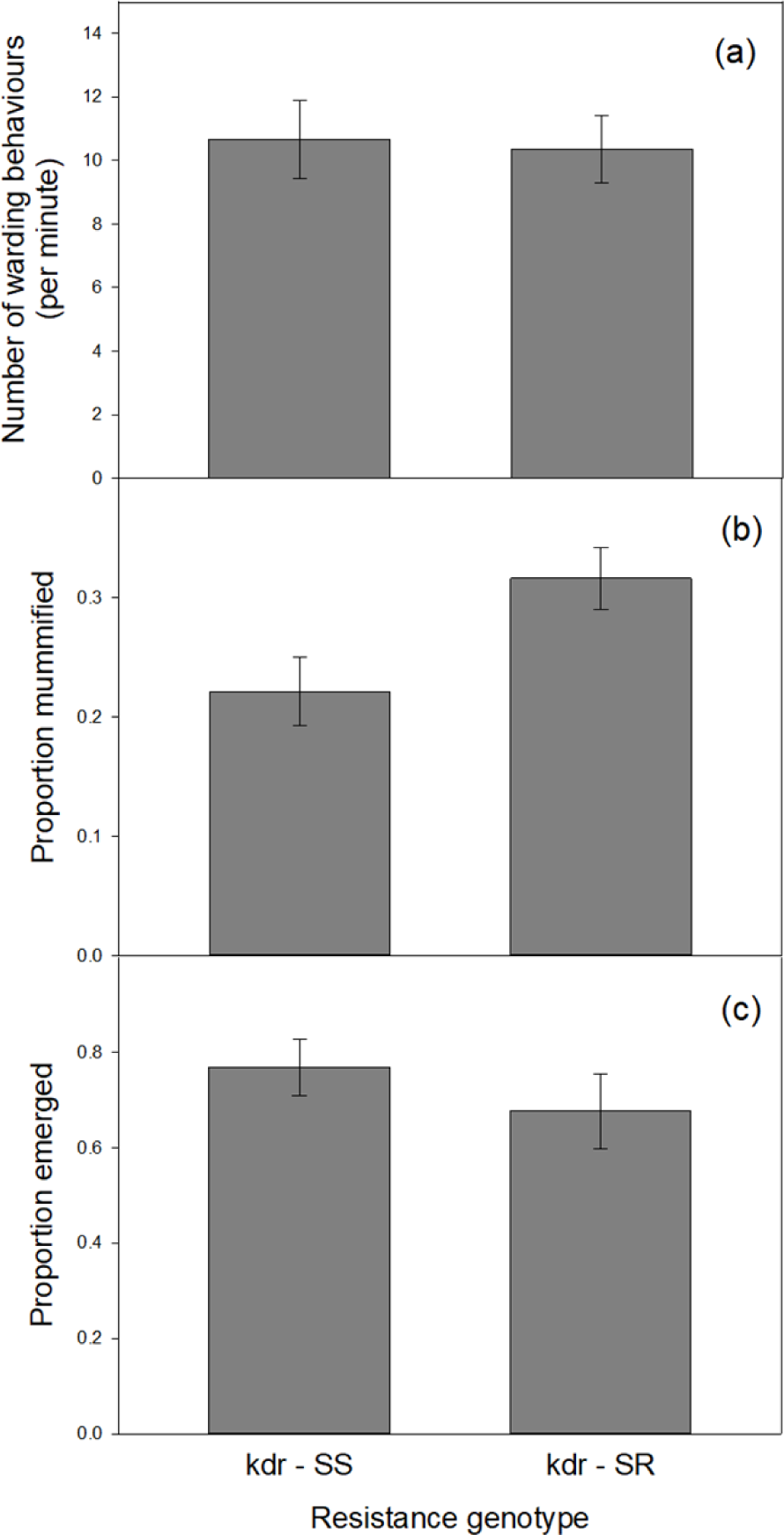
(a) Mean number of kicks/drops per minute carried out by the kdr-SR and kdr-SS clones when attacked by the parasitic wasp, *Aphidius ervi*. (b) The mean proportion of aphids mummified for both genotypes. (c) The mean proportion of wasps that successfully emerged from aphid mummies for both genotypes. Error bars represent ± individual SE for each category.

#### Mummification rate

Figure 3b illustrates the reduced proportion of mummified aphids of the kdr-SS clone compared with the kdr-SR aphids (F_1,6_ = 6.04, P < 0.049). The proportion of mummified kdr-SR aphids was 35% greater than that of the kdr-SS clone.

#### Aphidius ervi *emergence success*

Figure 3c illustrates that there was an decreased emergence rate of kdr-SR parasitoids from aphid mummies in comparison with the kdr-SS clones, but that this was not significantly different (F_1,6_ = 0.9, *P* < 0.38).

## Discussion

Throughout the course of the experiment, the pyrethroid resistant kdr-SR clone had a significantly lower tiller-level abundance than the pyrethroid susceptible kdr-SS clone. This may have been a consequence of the significantly reduced MRGR of the kdr-SR clone individuals compared with the kdr-SS clone. Reproductive rate is positively correlated with aphid size (Watt, 1979; Dixon and Dharma, 1980) and evidence shows that larger *Sitobion avenae* individuals have greater fecundity than smaller ones (Wratten, 1977). The lower population growth rate of the kdr-SR clone compared with the kdr-SS clone suggests either that the kdr-SR clones are less fecund than their counterparts and/or that they have increased mortality. Dixon (1970) demonstrated that smaller sycamore aphids (*Drepanosiphum platanoides*) have higher rates of mortality and that they are more likely to die before reproducing. A further possibility to explain the reduced kdr-SR abundance is therefore that they took longer to reproduce. Dixon and Wratten (1971) showed that smaller aphids take longer to produce their progeny. In their study, by the tenth day of adult life, large apterous aphids had produced approximately 60% of their offspring, whereas small apterous aphids had only produced 44%. The lower MRGR of the kdr-SR individuals may therefore have led to the increased time needed to produce all their progeny. This is supported by the observation that the kdr-SS population plateaued during the last week of the experiment, whereas the kdr-SR population was still increasing at the last count.

In addition to the slower population growth rate and reduced MRGR, the kdr-SR clone was also significantly more susceptible to parasitisation than the insecticide susceptible kdr-SS clone (31% vs 22% mummification rate, respectively). This is in agreement with Foster *et al*. (2007) who showed that insecticide resistant peach potato aphids (*Myzus persicae*) also had a greater rate of mummification compared with their insecticide susceptible counterparts. Foster *et al.* (2007) further went on to show that this was associated with reduced warding behaviour in the insecticide resistant clones. This study however failed demonstrate a difference in the ability of the two clones to exhibit behaviours intended to repel parasitoid attack. Upon first contact with the parasitoid both clones exhibited kicking or dropping behaviour approximately 10 times per minute. The explanation for the increased mummification rate of the kdr-SR clone therefore cannot lie with reduced warding behaviour and may possibly be due to reduced effectiveness of warding behaviour by the smaller kdr-SR clones.

Contrary to expectations there was no significant difference in parasitoid emergence success. The suitability of a host has been shown to affect parasitoid development (Harvey *et al.* 2014) and smaller hosts are less likely to provide the nutritional quality needed for parasitoids to develop and emerge (Desneux *et al.* 2009). *A. ervi* larvae require an intricate combination of endosymbionts and teratocytes provided by the host in order to grow exponentially within a mummy. Suboptimal teratocytes and endosymbionts provided by smaller hosts can drastically impair the physiology of parasitoid larvae (Pennacchio *et al*. 1999). The explanation for the unexpected lack of difference may lie with the relatively benign conditions found within the controlled environment, although this remains to be tested in a field situation.

The sudden appearance of the kdr mechanism in this SA3 *Sitobion avenae* clone appears to be a case of ‘forced evolution’, in which the development of the insecticide-resistant gene has led to numerous inhibitory, pleiotropic costs. Adaptations that evolve over a long period of time are likely to be more successful than rapid forced evolution and may not appear with these significant trade-offs. It may be that the fitness trade-offs acting against pesticide resistance have been intensified (McKenzie, 1996) due to the rapid kdr-mutation. As the kdr-SR aphids performed significantly less well than the kdr-SS clone in three of the five behavioural and physiological performance traits measured in this experiment, it is likely that this is the case.

This study suggests there is further potential to incorporate parasitoids into pest management schemes. The increased rate of mummification has shown that parasitoids can exploit trade-offs in the insecticide resistant *Sitobion avenae* clone which could possibly act to combat insecticide resistance. If the SA3 lineage acquires the ability to reproduce sexually, perhaps producing a kdr-RR genotype, it may exhibit an even greater level of immunological resistance. Should this be the case a strategy will be required to minimise the spread of this genotype and parasitoids would play a crucial part in this. Numerous studies have documented the success of parasitoids as components in agroecosystems (Starý, 1987). The alteration of cropping systems to favour natural enemies is often referred to as habitat modification and can operate at the plant, farm or landscape level (Gurr *et al*. 2004). It consists of diversifying crop ecosystems to make the environment more suitable for beneficial species. The addition of plant species that supply a resource, such as nectar or shelter, in an agro-ecosystem environment, has been proven to increase fecundity and longevity of parasitoids (Baggen and Gurr, 1998; Irvin *et al*. 2006). By making the environment more suitable to parasitoids, there is potential to reduce yield loss both directly and indirectly through the spread of aphid transmitted crop disease.

Furthermore, parasitoid behaviour can also be manipulated in order to maximise pest management. Ostensibly, the greatest difficulty in using parasitoids for biological control is synchronising the emergence of parasitoids with the beginning of aphid colonisation. If parasitoid emergence is not synchronous with aphid infestation, then aphid numbers become too large (Powell and Pickett, 2003). Parasitoids can be manipulated using aphid pheromones. Synthetic aphid sex pheromones have been shown to attract parasitoids (Powell *et al*. 1993) and exploring the potential of using such pheromones to manipulate female parasitoids with the aim of creating overwintering reservoirs in field margins, would be a valuable area of future research. This would allow parasitoids to swiftly recolonise crops in anticipation of aphid arrival in spring. In addition, plant volatiles, which are emitted when herbivorous arthropods feed on a plant, can be used by parasitoids as host location cues (Steinberg *et al*. 1993; Takabayashi *et al*. 1994). By producing crop varieties that release plant volatiles faster and in greater quantities, *A. ervi* would be able to locate aphids earlier (Powell and Pickett, 2003). In a field study Simpson *et al*. (2011) found that providing a combination of synthetic herbivore-induced plant volatiles and nectar plants improved the recruitment and residency of parasitoids and other beneficial arthropods.

Although there are many options available for using parasitoids as a biological control, the extensive use of pesticides will hamper such efforts in the future. Exposure to sub-lethal doses of the pyrethroid insecticide, lambda-cyhalothrin, have been proven to directly affect the ability of *A. ervi* to locate and oviposit on a host (Desneux *et al*. 2004). Furthermore, while the mummified shell can protect parasitoids from the effects of some insecticides throughout late larval, prepupal and pupal stages (Borgemeister *et al*. 1993; Jansen, 1996), other pesticides remain able to penetrate the mummified shell, subsequently killing the developing parasitoid (Hsieh and Allen, 1986; Krespi *et al*. 1991). A laboratory experiment conducted by Purcell and Granett (1985) found that the parasitoid *Trioxys pallidus* had a decreased emergence rate of 97% when sprayed with the organophosphate pesticide, azinphos-methyl. The authors found that 50% died within a day of emerging successfully, 39% died from ingesting toxic residue when cutting an emergence hole and 8% died within the mummy. Such studies put into question the effectiveness of parasitoids as a biological control when administered alongside pesticides.

However, numerous field studies have concluded that the effects of pesticides on parasitoids are significantly lower in comparison to laboratory bioassays (Obrtel, 1961; White *et al*. 1990). When compared to mummies sprayed with pesticide in a laboratory, mummies collected from crops in the field had a significantly lower mortality rate (Orbtel, 1961). The author provided two possible explanations for this variation. Firstly, pesticide residues sprayed in the field are more likely to experience weathering effects, such as photodecomposition, at a greater rate than laboratory conditions. These effects make them less toxic to emerging parasitoids. Secondly, mummies in the field receive less spray deposition than those in a concentrated laboratory environment. Therefore, various studies have described laboratory experiments as being a ‘worst-case scenario’ and highly unlikely to be representative of natural conditions (Longley, 1999).

